# Decoding neural responses with minimal information loss

**DOI:** 10.1101/273854

**Authors:** John A. Berkowitz, Tatyana O. Sharpee

## Abstract

Cortical tissue has a circuit motif termed the cortical column, which is thought to represent its basic computational unit but whose function remains unclear. Here we propose, and show quantitative evidence, that the cortical column performs computations necessary to decode incoming neural activity with minimal information loss. The cortical decoder achieves higher accuracy compared to simpler decoders found in invertebrate and subcortical circuits by incorporating specific recurrent network dynamics. This recurrent dynamics also makes it possible to choose between alternative stimulus categories. The structure of cortical decoder predicts quadratic dependence of cortex size relative to subcortical parts of the brain. We quantitatively verify this relationship using anatomical data across mammalian species. The results offer a new perspective on the evolution and computational function of cortical columns.

## Introduction

The mammalian cerebral cortex of mammals is a thin-layered tissue that appears to be assembled from a circuit motif termed the minicolumn, or column for short (Buxhoeveden, 2012). Each column spans the cortical layers and has stereotypic connections between cell types within and across layers. Columns can form ‘macro-columns’, which are groups of ∼100 minicolumns that are bound together by short range connections between columns. Even within macrocolumns, however, one can still discern the vertical structure corresponding to individual columns (Buxhoeveden, 2012). Although there are quantitative variations in column parameters across species, across brain regions, and even within a given macro-column, the main features of this circuit motif are quite universal. Columns are found in both sensory and motor areas of the brain (Harris and Shepherd, 2015), and analogous circuit motifs have also been found in non-mammals, such as birds (Wang et al., 2010). These facts strongly suggest that this circuit motif performs fundamental element(s) of a computation that is needed independent of the stimulus modality. However, the complexity of connections, and the large number of cell types within the column (many of which remain to be strictly defined (Harris and Shepherd, 2015; Jiang et al., 2015; Luo et al., 2017)), have made it difficult to determine the algorithm implemented by cortical columns.

To gain insights into the computations performed by cortical columns, one can begin by analyzing, from first-principles, possible strategies for representing stimuli with neural responses in ways that allow for their accurate decoding, ideally with minimal loss of information. The colloquial term “information” used here can be quantitatively defined using tools from information theory (Cover and Thomas, 1991). In the context of this work, by information we mean the mutual Shannon information (Cover and Thomas, 1991) between stimuli and neural responses. When considered in small time intervals, neurons respond to stimuli by producing all-or-nothing events in the voltage traces across their membranes termed “spikes”. Unfortunately, unlike the genetic code where it is known how to parse DNA sequence to determine amino acid sequence, it remains unclear how to parse sequences of spikes over time to determine which stimuli they represent (Srivastava et al., 2017; Theunissen and Miller, 1995). Similarly, it is a matter of debate for how to combine spikes from neurons tuned to different stimulus features. A simple weighted average of responses across neurons has been shown to discard substantial amounts of information (Osborne et al., 2008; Reich et al., 2001), indicating that more complex codes are used in the brain. We will discuss how insights into these problems can be gained by searching for the code that allows for decoding with minimal loss of the information contained in neural responses. Further, we present evidence one version of this code is implemented by cortical columns.

## Results

### A code with information-preserving statistic

We begin by describing how a stimulus can be represented by neural responses in a way that these neural responses can be decoded without loss of information. For ease of exposition, we will first consider the case where, once the stimulus is specified, noise in neural responses is independent across neurons and across different time bins (we will show that the main result holds even when these constraints are removed). Further, we will initially analyze stimulus representations where the responses of any single neuron depend only on one stimulus dimension (this dimension corresponds to neuron’s receptive field (RF), as we mathematically describe below). Later we will see that expanding representations to allow dependence of neural responses on multiple stimulus dimensions is one of the key distinguishing features of the cortical column decoder, compared to a simpler decoder.

With these assumptions, the neural responses can be described using the so-called linear-nonlinear (LN) model (Schwartz et al., 2006). In this model, the probability of observing a spike depends on the strength of the relevant stimulus component (Figure 1). The nonlinear dependence of the neuronal spike rate on the primary stimulus component can often be well approximated by a saturating (logistic) function that has two parameters: a threshold *α* and a steepness *β*. We note that neural responses can often be also equivalently described using tuning curves (Abbott and Dayan, 1999; Georgopoulos et al., 1986; Hohl et al., 2013; Osborne et al., 2008; Shamir, 2014). Tuning curves specify how neural response rates change when stimuli deviate from their optimal settings. There is a one-to-one relationship between the parameters of the tuning curves and those of the saturating nonlinearity in the LN model (Supplementary Text 1). However, the LN formulation makes it possible to mathematically derive the vector quantity 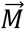 that is guaranteed to capture all the information provided by the responses of the neural population (cf. Supplementary Text 2). This quantity is constructed as:

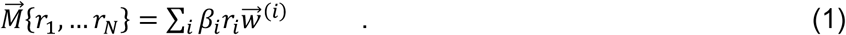

**Fig. 1.**
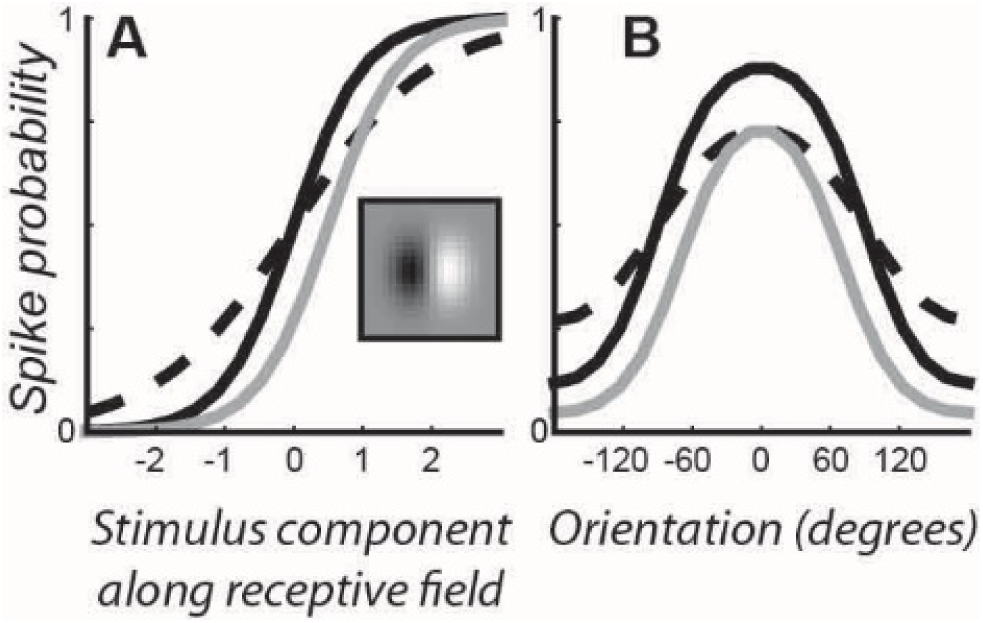
The relationship between receptive field (RF) and tuning curve descriptions of the neural response. (**A**) Three representative model nonlinearities that describe neural response as a saturating function of stimulus component along RF. Black and dashed lines have the same midpoints *α* but different steepness values *β*. Black and gray lines have same β values but different *α* values. Inset shows an example orientation selective RF. (**B**) Corresponding tuning curves from (**A**) but as a stimulus function of angle.

In this expression, *r*_*i*_ denotes the number of spikes produced by the *i*th neuron during the time interval of interest, *β*_*i*_ is the steepness value of the *i*th neuron nonlinearity, and 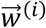 is the preferred stimulus for this neuron, also known as neuron’s RF, normalized to have unit contrast (mathematically, 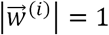). We will refer to 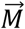 as the information-preserving population vector. This vector generalizes the standard population vector (Georgopoulos et al., 1986; Hohl et al., 2013; Salinas and Abbott, 1994; Shamir, 2014) by taking into account steepness parameters *β*_*i*_. In the context of the LN model, steeper nonlinearities indicate more reliable neural responses. Therefore, it is perhaps intuitive that the responses of more reliable neurons should be weighted more strongly within the population average.

Taking into account steepness parameters fully addresses previous concerns regarding the standard population vector, namely that averaging responses of similarly tuned neurons can lead to substantial information loss (Osborne et al., 2008; Reich et al., 2001). In Figure 2, we show that our information-preserving version of the population vector captures all the information available in neural responses, regardless of whether the neurons in the population are tuned to the same (Fig. 2A) or different (Fig. 2B) features of the stimulus. In contrast, the standard population vector does not capture all of the information when the population contains neurons tuned to the same features of the stimulus (or features with opposite polarity) using nonlinearities with different steepness values.

**Fig 2.**
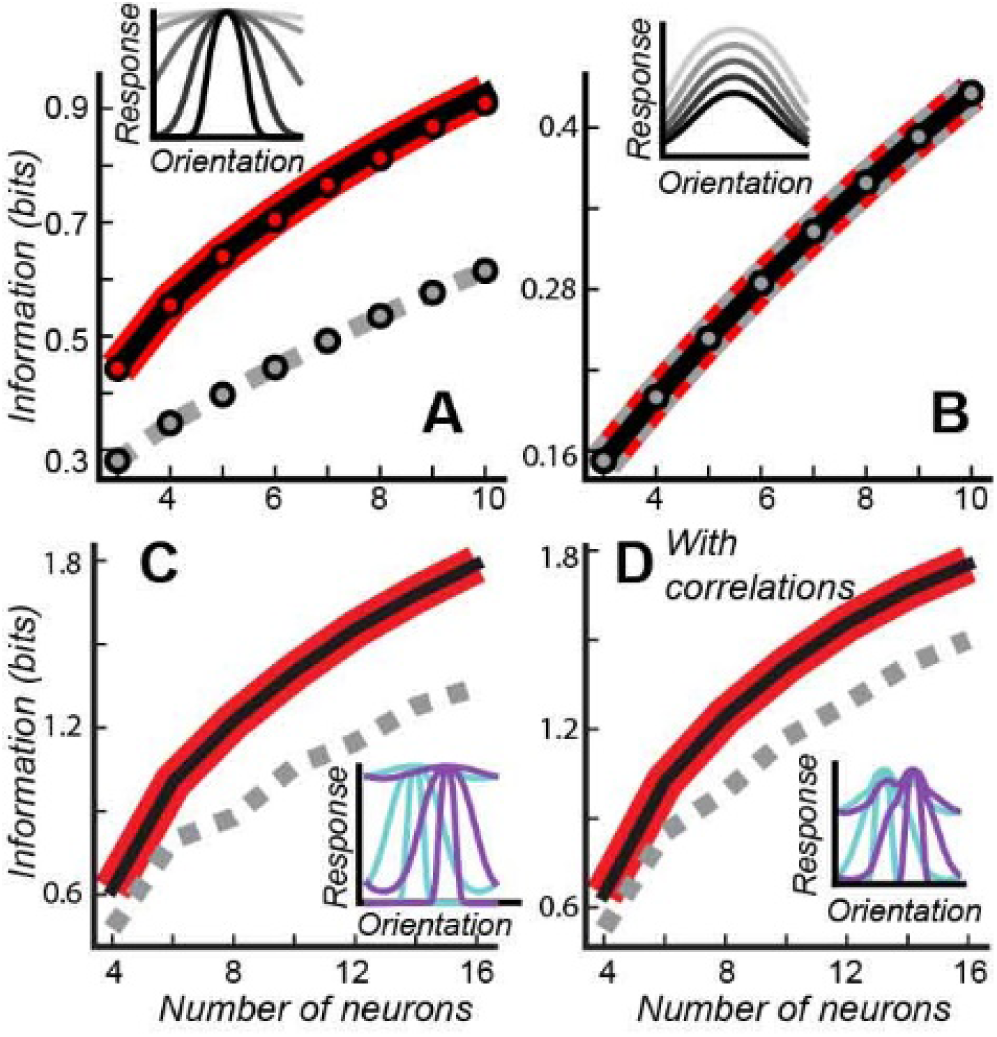
The information-preserving vector captures all information from diverse neural populations and with correlated variability across neurons. Neural populations tuned to the same (**A**, **B**) or different (**C**, **D**) preferred stimuli. In (**A**) differences in neural tuning curves are due to differences in steepness values, whereas in (**B**) they are due to differences in thresholds. Lines formed by dots show the information values obtained by binning response variables. Dotted lines overlap with solid curves. (**D**) same as (**C**) but with noise correlations. Insets show example population tuning curves for *n* = 6. In all panels, we compare information transmitted by a population response (black line) with information transmitted by the information-preserving population vector (red) and the standard population vector (dashed gray).

It is worth noting that some differences in neural tuning curves are irrelevant for capturing all the information contained in neural responses. For example, although differences in thresholds lead to differences in tuning curves, according to the information-preserving expression (1), responses of neurons with different thresholds can still be averaged without losing information. Figure 2C shows that both the standard and information-preserving population vectors capture all the information in model neural population without taking threshold differences into account. These analyses demonstrate that analyses in terms of saturating nonlinearities are more revealing than those based on the tuning curves.

The information-preserving population vector also works in the presence of correlated variability across neurons (the so-called noise correlation reviewed in (Averbeck et al., 2006; Shamir, 2014)). Although noise correlations may affect the overall information provided by the neural responses (Abbott and Dayan, 1999; Ecker et al., 2011; Moreno-Bote et al., 2014; Shamir, 2014; Shamir and Sompolinsky, 2004, 2006; Zohary et al., 1994), the information-preserving population vector continues to capture all of the information that is available in the neural responses (Fig. 2D). This result holds true for noise correlations that differ across pairs of neurons (e.g. according to differences in RFs) as long as noise correlations do not change with the stimulus, as is often observed experimentally(Huang and Lisberger, 2009). In Figure 2 we show the results of model simulations with noise correlations and provide a detailed derivation in the Supplementary Text S3. We also note that in the presence of noise correlations, the nonlinearities of individual neurons may deviate from the logistic function but the information-preserving property still holds as long as the population response can be written in the exponential form (see Eq. S24 in the supplement).

### Capturing information in cortical responses

Analyses of the model neural populations show that the information contained in the responses of model neurons conforming to the LN model can be fully captured by a version of the population vector that is modified in a specific way. Although the LN model has been successfully used to describe neural responses in a number of brain circuits (Schwartz et al., 2006), to the extent to which real neural responses deviate from the model assumptions, some information loss will occur. To determine the magnitude of these effects, we tested how the information-preserving population vector performs on the responses of neurons in the primary visual cortex (V1) that were elicited by natural stimuli (Sharpee et al., 2006). For each neuron, we estimated its preferred orientation and nonlinearity. Nonlinearities were fit to the neural responses using logistic regression to find the steepness parameters *β*. Based on the estimates of β values, we compared the full amount of information provided by the responses of these neurons with the information provided by the standard and information-preserving population vectors. To account for experimental uncertainties in the orientation value, we used a coarse-grained set of orientations that took into account error-bars (see Materials and Methods). We find that just like in the model neural populations, the information-preserving population vector (but not the standard population vector) captured all the information (Figure 3) provided by the responses of simultaneously recorded neurons.

**Fig. 3.**
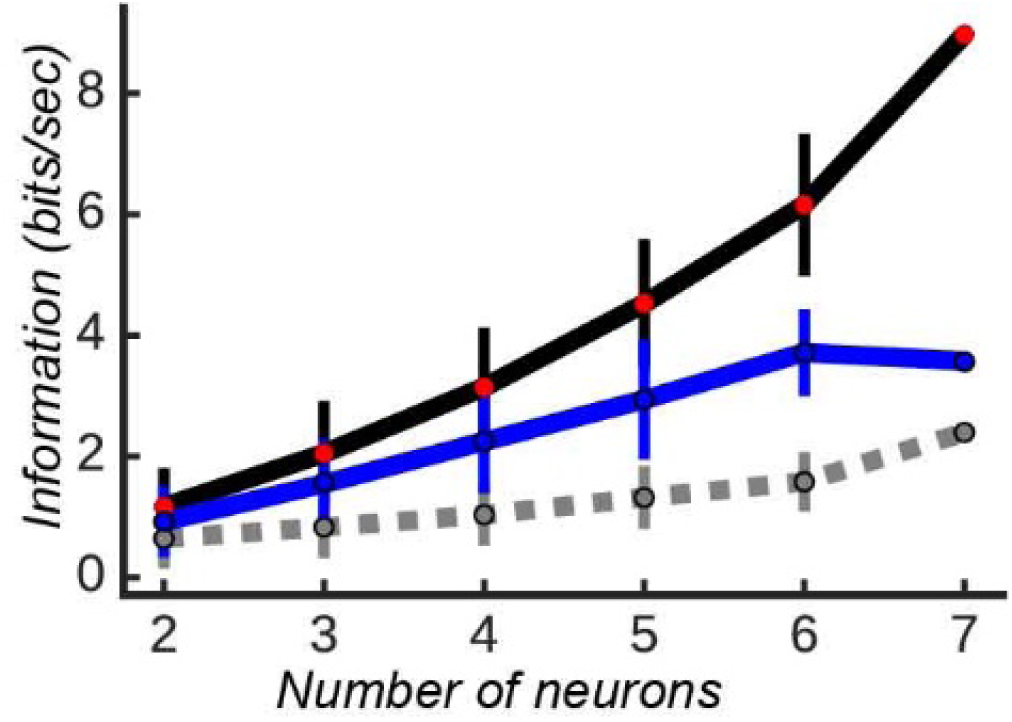
The information-preserving population vector captures all the information provided by responses of simultaneously recorded V1 neurons. Curves for the information provided by neural responses (black line) and that captured by the information-preserving population vector (red circles) overlap. The standard population vector (blue) and population count (grey) provide smaller amounts of information. Error bars are standard deviations.

### Decoding algorithm

We now show how signals can be decoded from the information-preserving population vector. The value of this vector varies continuously with the stimulus, because different stimuli evoke different responses *r*_*i*_. The expected value of the information-preserving vector depends on the stimulus 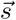 as:

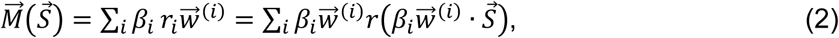

where *r(x)* is nonlinear response function and parameters 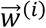 and *β*_*i*_ are same as before in Eq. (1) (see Text S3 for derivation). To understand the properties of this mapping, we can build on research in the area of RF estimation (Schwartz et al., 2006), where this mapping uses responses of a single neuron to many different stimuli to estimate that neuron’s RF. Here we use RFs of many neurons and their responses to a single stimulus to estimate that stimulus. Based on studies of RF estimation, we can state that the information-preserving population vector will be aligned with the stimulus multiplied by the covariance matrix *C* of RF components across the neural population under certain statistical conditions (Supplementary Text S4 for a derivation and discussion of deviations when these conditions are not met).

Crucially, using a standard population vector expression 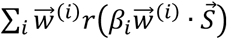 (or any expression where RFs are not scaled by the same factor *β*_*i*_ from the spike rate nonlinearity) introduces estimation biases. This is shown in simulations of Figure 4A and mathematically in Text S4. These biases arise as soon as RF components have non-equal variance in different stimulus directions. Such is the case, for example, in V1 where more RFs align with horizontal and vertical orientation than with oblique angles (Dragoi et al., 2001). We have also verified these results using neural data by reconstructing segments of natural movies using either the information-preserving or standard population vector, as well as a population vector with random *β*_*i*_ factors. The information-preserving vector produced significantly more accurate reconstructions than either of the two alternatives (p< 10^−39^ t-test, Fig. 4B). Further, the decoder maintains much of its accuracy even if the estimates of *β* factors are imprecise. For example, we used a decoder where the *β* factors were multiplied by a random value between 0 and 1; such a decoder yielded correlation coefficients that, on average, were more than half of the correlation values provided by the true information-preserving population vector. Thus, even partial knowledge of *β* factors can result in substantial improvements in decoding accuracy.

**Fig. 4.**
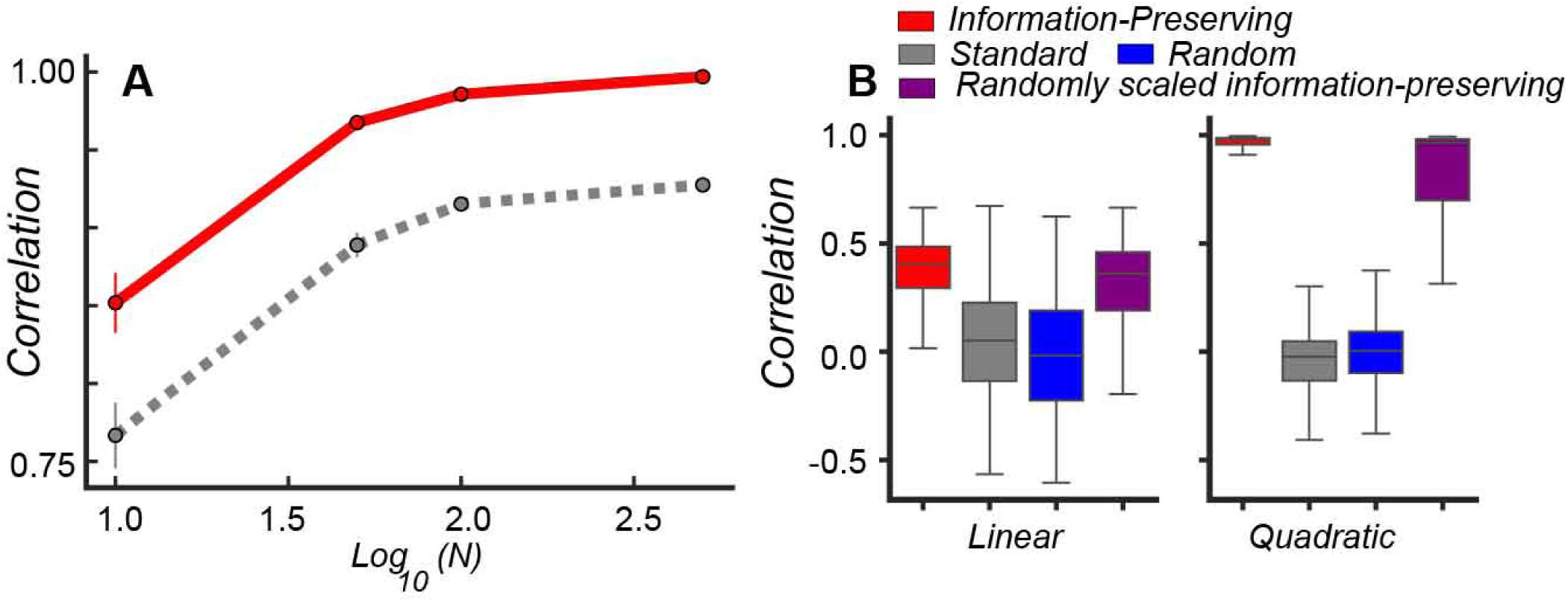
**(A)** Average correlation between stimuli and their reconstructions based on either the information-preserving (red) or standard population vectors (grey). The information-preserving decoder converges to an unbiased estimate of stimulus (red line), but not the standard population vector (dashed grey). Error bars are standard error of the mean across realizations of population RFs. (**B**) Test of decoders based on V1 data (86 neurons) from (Sharpee et al., 2006). The reconstructions do not require that all of the neurons are recorded simultaneously. Therefore, we could use an expanded set of 86 V1 neurons for which the responses to the same set of natural stimuli were available. The box and whisker plots show median and interquartile range (IQR); whiskers are 1.5 of IQR. Correlation values here reflect measurements for individual stimuli, not averaged across the set of stimuli.

### Feedforward neural network decoder

In terms of biological implementation, stimulus decoding based on the information-preserving vector can be implemented using a three-layer feedforward neural network, as illustrated in Figure 5A. Units within the first layer encode stimulus components; the second layer provides a representation according to Eq. (1); units within the third layer units represent reconstructed stimulus values according to Eq. (2). The key aspect of this decoding scheme is the incoming connections to each second layer unit should be proportional to outgoing connections. A version of this decoder was shown to accurately reflect the synaptic and network mechanisms of the leech nervous system (Lewis and Kristan, 1998). Similar networks operate in subcortical areas in mammals (Joshua and Lisberger, **2015**), and can account for initial stages of olfactory processing (Zhang and Sharpee, 2016) for odor mixtures. We note that this decoder has compressive nonlinearities (illustrated in Figure S1), making it different from the optimal linear decoder proposed previously (Salinas and Abbott, 1994).

**Fig. 5.**
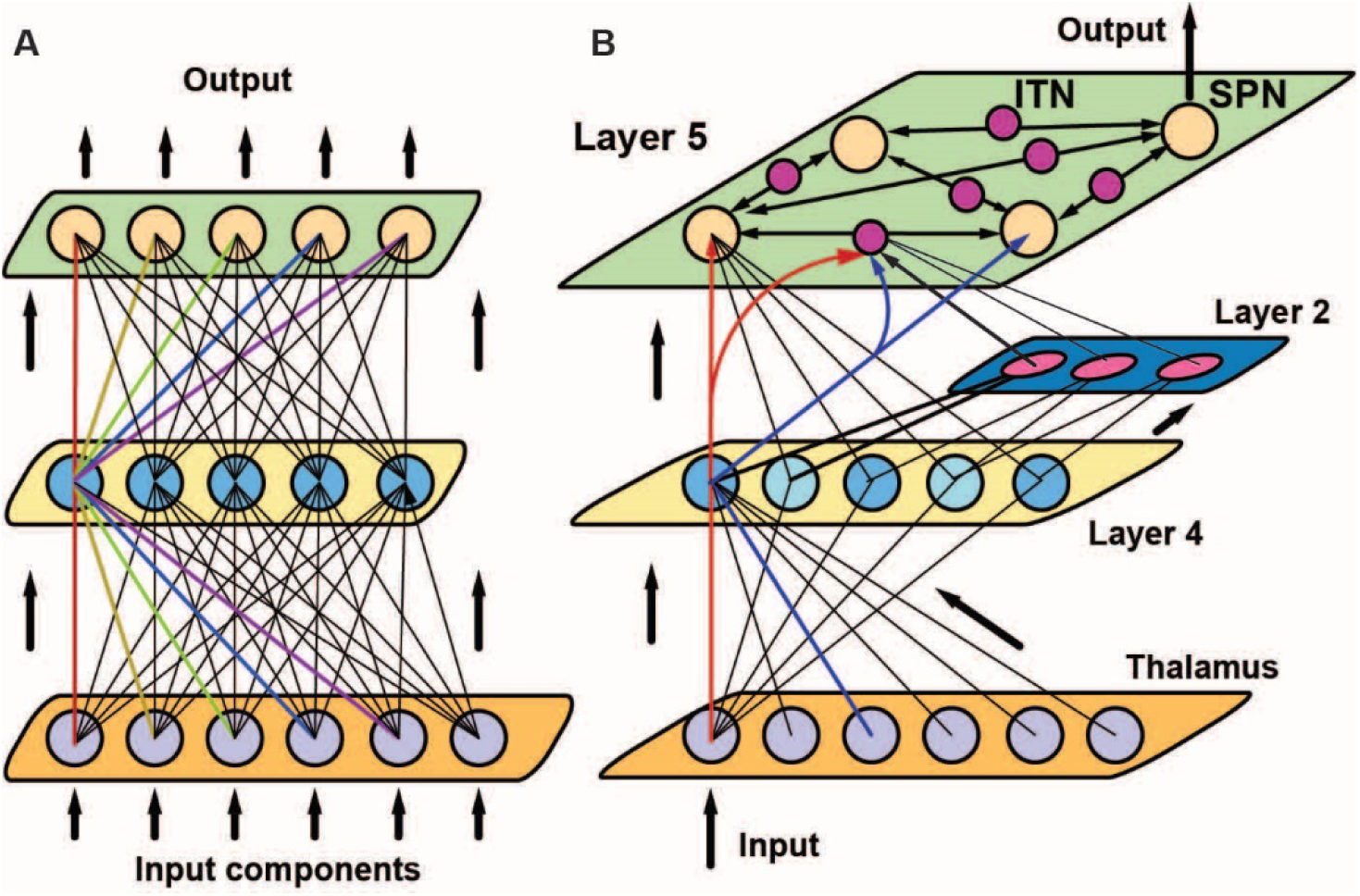
Schematic of a feedforward (A) and quadratic (B) decoder. (**A**) A feedforward network can implement the decoder in the original input space. Color marks connections of the same strength. (**B**) Quadratic decoding adds recurrent computation in layer 5. Input from layer 2 complex cells to layer 5 intratelecephalic neurons (ITNs) sets the recurrent weights between the output layer 5 subcerebral projection neurons (SPNs). For clarity, only a subset of connections is shown.

### Decoding of ambiguous stimuli

We now show how the decoding scheme can be generalized to increase its accuracy and to allow neural circuits to deliberate between alternative, mutually exclusive interpretations of ambiguous stimuli. This is an important problem because sensory perception is in general ill-defined, meaning that different stimuli can give rise to similar patterns of neural responses. The decoder discussed so far (e.g., Figure 4 and 5A) does not solve this problem because for each stimulus it produces a single interpretation. To allow for multiple interpretations, one can expand the stimulus representations quadratically by including all pairwise products *s*_*i*_*s*_*j*_ between original *s*_*i*_ components. That is, if the original stimulus has *D* components, after expansion, the stimulus will be represented by the original *D* components plus an additional *(D*^2^*+D)/2* pairwise components. Working in this expanded space, one can construct the information-preserving population vector according to Eq. (1) as before. The information-preserving vector now has a linear part *M*_*i*_ and a quadratic part *M*_*ij*_ (see Supplemental Text S5 for details). To decode a single stimulus pattern from these two parts, we need to find a pattern *s*_*i*_ that approximates the matrix *M*_*ij*_ as best as possible in the form of *s*_*i*_*s*_*j*_. Mathematically, this operation corresponds to finding the leading mode of matrix *M*_*ij*_. A key property of this transformation is that multiple modes can potentially provide similar contributions to the matrix. This situation corresponds to ambiguous stimuli, with each mode describing alternative representations. The conflict between modes can be resolved by waiting for additional evidence to favor a specific representation and/or by incorporating evidence from larger scales. Thus, decoding with quadratic stimulus expansion represents a conceptual advance compared to the decoding in the original input space. Of course, to allow for the possibility of multiple modes, the original stimulus should be multidimensional with *D>1.* It also should be noted that the purely quadratic decoder based on *M*_*ij*_ does not determine stimulus polarity. The stimulus polarity is determined by comparing the sign of the estimated stimulus with the linear part *M*_*i*_. Thus, both parts of the information-preserving population vector are needed to ensure complete stimulus reconstruction.

We tested the accuracy of this quadratic decoding algorithm on recoded V1 responses(Sharpee et al., 2006). We found that it produces improved reconstructions compared to those made without quadratic stimulus expansion (p<10^−28^, Fig. 4D). Furthermore, just like in the original stimulus space, a reconstruction based on the information-preserving population vector is significantly better than those based on the standard population vector or population vector computed with randomly selected *β*_*i*_ values (p<10^−52^). Finally, similar to decoding without quadratic stimulus expansion, the quadratic decoder is robust to noise in the estimation of *β*_*i*_ factors. In Fig. 4b we show that decoding using *β*_*i*_ factors that have been multiplied by a random number from 0 to 1 produces accurate stimulus reconstructions. We observe that quadratic stimulus decoding is even more robust to this perturbation than decoding performed in the original stimulus space (Fig. 4B).

### Recurrent neural network decoder

Given the computational benefits of decoding with quadratic stimulus expansion, how can it be implemented in neural circuits? We now discuss how each of the steps of this algorithm maps onto computations performed by the cortical column. First, one needs neurons with RFs in the quadratically expanded input space (Fitzgerald et al., 2011). When analyzed in the original input space, these neurons would have quadratic nonlinearities (possibly also with a non-zero linear component). In V1, such neurons are known as complex cells (Movshon et al., 1978). However, they are also found in upper layers of the primary auditory cortex (Atencio et al., 2009). The responses of these neurons can be obtained by adding the responses of simple cells from layer 4 that are selective for stimuli of opposite polarity. This corresponds to the classic model of complex cells responses (Movshon et al., 1978), and which is also consistent with strong projections from layer 4 simple cells to layer2 complex cells (Harris and Mrsic-Flogel, 2013; Harris and Shepherd, 2015).

Next, the information-preserving population vector must be computed both in the original space and in the quadratically expanded space. In the original input space this computation can be done using the same feedforward procedure as before, in this case by pooling the responses of simple cells according to Eq. (2) 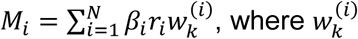 stands for the *k*th component of *i*th neuron RF, and *N* is the number of simple cells. The quadratic part of the information-preserving population vector *M*_*kn*_ can be estimated by pooling the responses of complex cells *c*_*i*_ to form 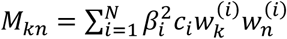. The weights 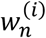 can be provided by copies of connections from the corresponding simple cells.

The final step is to find the dominant mode of the matrix represented by quantities *M*_*kn*_, choosing its polarity based on *M*_*i*_ values. These two operations can be simultaneously computed by a recurrent network that receives quantities *M*_*i*_ as inputs to its *i*th neuron and connections between neurons *k* and *n* in the network are set to *M*_*kn*_. In the presence of gain normalization (Carandini and Heeger, 2011), the activity of this network will converge to the dominant mode of matrix, implementing the so-called power method.

Based on the circuitry of cortical columns (Harris and Mrsic-Flogel, 2013), one can identify the output recurrent network with that of layer 5 subcerebral projection neurons (SPN) as well as layer 6 cortical thalamic neurons. These neurons receive connections from layer 4 that necessary to compute input values *M*_*i*_ A key aspect of this recurrent optimization is that connection strengths *M*_*kn*_ must vary with the stimulus. The layer 5 contains a population of cells termed intratelecephalic neurons (ITNs) that project to layer 5 SPNs and can modulate connections between SPNs in a stimulus-dependent manner. The ITNs receive signals from layer 2 necessary to compute *M*_*kn*_ values. Thus, the cortical column has all of the required components to implement quadratic decoding.

### Predicted scaling relationships

If quadratic decoding is indeed the algorithm that is being implemented by cortical columns, then this yields a number of quantitative predictions concerning for the distribution of cell types across layers, and how the size of the cortex should scale with the number of subcortical inputs. Here we review these predictions in turn.

The first prediction is that the number of ITNs in layer 5 should equal to the square of the number of output SPNs in layer 5. There is evidence in the mouse motor cortex that this is the case. Anatomical images from indicate that there are 8-9 corticospinal neurons and 60-80 ITNs in each minicolumn (Oswald et al., 2013) The second prediction describes how the total number of neurons in a column should scale with the dimensionality of the signal. Specifically, a minicolumn that processes *D*-dimensional signals should have *∼D*^2^ *+ 𝒳*^*D*^ neurons. The quadratic term arises from the need to implement stimulus-dependent recurrent weights between the output neurons. The linear term includes the output neurons as well as neurons from the intermediate representations, such as simple and complex cells, whose number should be proportional to *D*. For a piece of cortical tissue with *N*_columns_, the number of neurons will be *N* cortex = *N* columns (*D*^*2*^ + *𝒳*^*D*^) The corresponding number of subcortical input neurons *N*_t_ = *N*_columns_ *D*. Combining these two relations yields the following prediction

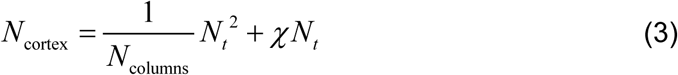

In Figure 6, we show that the predicted quadratic function accounts well for the differences in the number of cortical and subcortical neurons (from the diencephalon, brainstem, and basal ganglia) across 18 mammalian species (Herculano-Houzel, 2009). We fit data across primate species separately because they are known to have scaling exponents that are different from other mammals in a number of ways (Herculano-Houzel, 2009). For rodents/insectivores, the fit yields: *N*=_columns_ (2.7 ± 0.9)108, 𝒳 = 1.75 ± 0.16; for primates the fit yields: *N*_columns_ = (1.6 ± 0.4)108, *𝒳* = 11.7 ±1.5. Although the parameter values *N*_columns_ are obtained merely by fitting the data, without any constraints, they quantitatively match the current estimate of 2 ×10^8^ for the number of minicolumns in the brain (Sporns et al.). The same estimate can also be obtained by dividing the total number of cortical neurons (Herculano-Houzel, 2009) by the estimated number of neurons per minicolumn (Buxhoeveden, 2012). Further, one can the use these parameters to derive estimates for the microscopic parameters of individual columns. Combining the values for the parameter *𝒳* with the estimated number of ∼100 neurons per minicolumn (Buxhoeveden, 2012) yields estimates for the number of output neurons *D* of ∼9 for rodents and ∼6 for primates, both of which agree with experimental values for these species (Oswald et al., 2013; Peters and Sethares, 1996). Thus, the scaling rules that follow from the structure of the quadratic decoder are supported by a diverse sets of quantitative predictions across nine orders of magnitude in neuron numbers (from 10 to 10^10^).

**Fig. 6.**
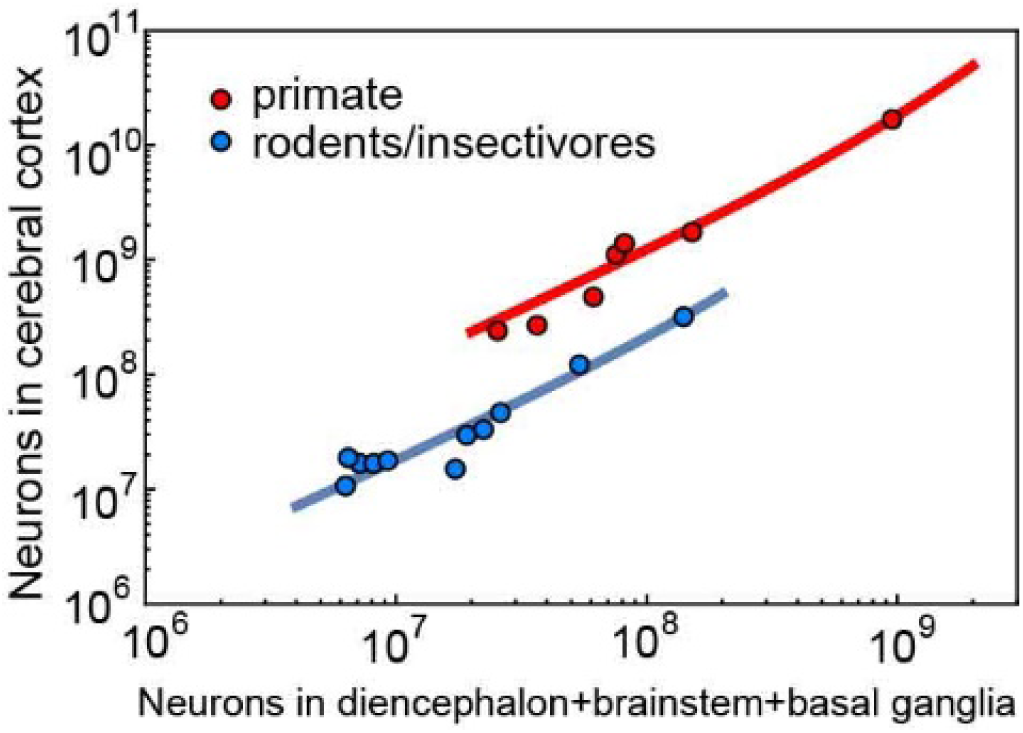
Quadratic decoding explains quadratic scaling of cortex size across species. The number of cortical neurons as a function of to the total number of neurons in brain stem, basal ganglia and the diencephalon, which includes the thalamus. Data from (Herculano-Houzel, 2009). Solid lines represent fits using Eq. (3).

## Discussion

In this work we started from first principles of information theory to find a set-up where stimuli can be encoded into neural responses in such a way as to enable the decoding of these responses with minimal loss of information. For this set-up, there a simple vector quantity that captures all of the information contained in neural responses. Based on this quantity, one can build two kinds of decoding algorithms. The first algorithm uses primarily feedforward operations of the kind found in invertebrate (Lewis and Kristan, 1998) and mammalian subcortical circuits (Joshua and Lisberger, 2015). Of course, subcortical circuits compute more sophisticated computations than just stimulus reconstruction. We analyzed stimulus reconstruction just as an example of a computation that neural circuits may perform.

A second more sophisticated way to perform decoding is to quadratically expand the stimulus space and then to use recurrent optimization to invert the transformation. We argue that the second decoding algorithm, which we tern quadratic decoding, is what cortical columns compute to deliver increased accuracy, as well as the ability to disambiguate between alternative stimulus interpretations. Quadratic decoding brings together many disparate properties of cortical processing as parts of a single computation. For example, the decoder requires synaptic weights to be zero on average for some neural populations (see Supplementary Text S4). This corresponds to the so-called balanced regime where inhibition and excitation on average cancel each other (van Vreeswijk and Sompolinsky, 1996). Furthermore, because recurrent networks can become unstable (Sompolinsky et al., 1988), a specific gain control is needed to reduce effective recurrent connections within layer 5 when necessary. The observed inhibitory gain control (Sompolinsky et al., 1988) that layer 5 exerts through layer 6 back to layers 4 and 2 can fulfill this role (Olsen et al., 2012). Some cortical areas, such as the olfactory system as well as the hippocampus are missing layers 2 and 4. These areas, therefore, likely lack this form of gain control, which would explain why they often serve as seizure origination points.

The structure of the quadratic decoder reveals new constraints on mammalian brain evolution. Two separate factors drive brain size expansion within and across mammalian species orders. The first factor controls brain expansion within orders, e.g. within primates. This factor represents the average dimensionality of inputs processed by each column. Among primates, humans have the largest value. The second factor *𝒳* controls brain expansion across orders, such as between rodents and primates. It represents the weighted number of neuronal types per input dimension, weighted by the number of neurons in each type. Because of this weighting and because excitatory neurons comprise a majority of neurons (Jiang et al., 2015), the factor *𝒳* mainly reflects diversity of excitatory neuronal types. Taking this into account, the derived value for rodents is consistent with current estimates for the number of cortical excitatory cell types (Jiang et al., 2015; Markram et al., 2015). Further, a large increase in this factor from rodents to primates is consistent with observations that excitatory neuronal types are less conserved between rodents and primates than inhibitory types. A larger number of cell types encoding signals along each dimension increases the accuracy with which each signal can be encoded, as has been demonstrated in the retina (Kastner et al., 2015). At the same time, primate minicolumns process signals of smaller dimensionality than rodent minicolumns. This allows for finer sampling and reduces the number of competing modes for each quadratic decoder. These analyses highlight different axes that evolution can manipulate to acha accurate decoding of complex stimuli.

## Author Contribution

JB derived information-preserving property, TS derived decoding algorithms, their implementation in neural circuits, the mapping onto cortical circuitry and the scaling relationships. Both authors analyzed the data and wrote the paper.

## Acknowledgments

We thank Vicki Lundblad for comments on the manuscript. This research was supported by the Rose Hill Foundation, the National Science Foundation (NSF) award numbers IIS-1254123, IIS-1724421, and IOS-1556388.

## Materials and Methods

### Estimating Mutual Information from V1 Recordings

We represent the responses of a set of *N* simultaneously recorded V1 cells to a stimulus binned into *T* segments and repeated *K* times as a tensor *D* of shape (*T, K, N*). *T* is typically 330, corresponding to time bins of 30 milliseconds. *D*_*tij*_ represents the number of times neuron *j* fired in response to stimulus *t* on repeat *i*, and can be any nonnegative integer.

### Converting Data to Binary Words

Let *v*_*max*_ be the largest value in *D*; the maximum number of spikes over all neurons, stimuli segments, and repeats. We form a binary tensor 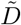 of shape (*v*_*max*_ × *T, K, N*) by resampling each slice Dt · into a sub tensor 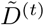 of shape (*v*_*max*_, *K, N*) according to the following algorithm:

1. For a given value of *i* and *j* we let *n* = *D*_*tij*_. We sample without replacement a set 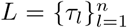 of *n* indices from the integers {1, …, *v*_*max*_*}*
2. We set 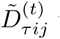 to 1 if τ *∈ L*, and to 0 if not.
3. Steps 1 and 2 are repeated for all *i* and *j*.

After all the time slices of D are resampled, the set 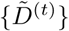 of binary tensors are concatenated to form a binary data tensor 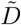 of shape (*v*_*max*_ · *T, K, N*). Each row of 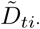 corresponds to a sample of {rj}. We note that the samples described by 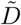 correspond to time bins of length 30/*v*_*max*_ milliseconds.

In Figure 3 we include only sets of neurons recorded simultaneously, and all subsets. Thus, a set of 4 simultaneously recorded neurons yields one set of size 4, 4 sets of size 3, and 6 sets of size 2.

### Estimating parameters of the linear-nonlinear models

To compute population vectors and the information captured by them we need estimates of the linear-nonlinear (LN) model parameters 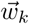 and *βk* for every neuron. In order to estimate *βk*, we fit the response rate of the *k*th neuron evoked by stimulus 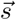 (averaged across the repeated presentation of this stimulus) using a logistic function:

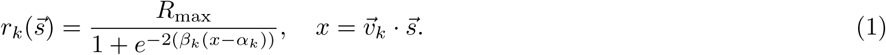

In this expression 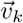 is estimated as the maximally informative dimension for the neuron [Sharpee et al., 2006]. The parameters *R*_max_, *α*_*k*_, and *β*_*k*_ are fit by minimizing the mean square error between 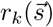 and experimentally measured firing rates.

To account for non-monotonic response functions, we also fit neural response functions using logistic function applied to a quadratic function of the stimulus. Specifically:

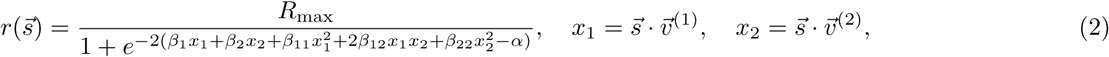

where 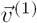 and 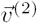 are two RF components estimated as the first and second maximally informative dimensions for the neuron [Sharpee et al., 2006]. These fits were used with quadratic decoding described in Text S5.

### Information captured by population vectors

For the analysis in Figure 3 we used 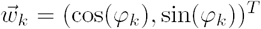, where *ϕ*_*k*_ are preferred orientation values estimated from the MID vector 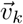 for each neuron computed in [Sharpee et al., 2006, Sharpee et al., 2008], along with their standard deviations Δ*ϕ*_*k*_. Additionally, in this figure we plot the information computed under a coarse grained realization of orientation values *ϕ*_*k*_ to take into account experimental errorbars Δ*ϕ*_*k*_ accociated with them. The coarse graining is based on the following measure of distinguishability between orientation values for neurons *i* and *j*:

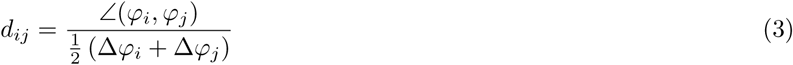

using the following procedure:

1. Find the pair of neurons (or subpopulations if multiple neurons have the exact same *φ*) with the smallest value of *d*_*ij*_.
2. Compute 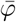 as the weighted angular average of all φ for the set of neurons in step 1, with weights given by 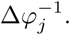 Similary compute the average value of Δ*φ*.
3. For all neurons in the set found in step 1, replace φ with 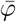 and *Δφ* with its average.
4. Repeat steps 1-3 until no pair of neurons with distinct *φ* have *d*_*ij*_ *<* 1.

### Adjusting for finite sample effects

We now describe how we estimated the information transmitted by a set of neural responses {rk} about a set of stimulus samples 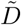 The process is the same for estimating the information transmitted by the standard and information-preserving population vectors, as they are also discrete random variables of known cardinality. Our information estimate is the finite sample approximation of Shannon’s Mutual Information

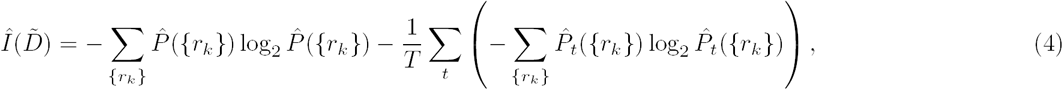

where 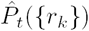 is the empirical probability of the population response equalling {*rk*} at time bin t, computed across repeats. The marginal distribution 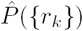 is simply the average of 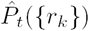 across time bins:

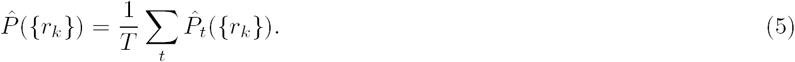

Because 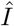 is a biased estimate of the true mutual information for finite samples [Treves and Panzeri, 1995, Strong et al., 1998],we corrected for finite sample effects by subsampling 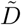 using the approach [Treves and Panzeri, 1995]. Specifically, we computed 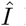 using a fraction *f* of the repeats, for *f ∈ {*1.0, 0.95, 0.90, 0.85*}*, sampling repeats without replacement. We performed this subsampling ten times for each value of *f*. We perform linear regression on the values of 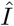 vs *f*^−1^ and extrapolate to *f*^−1^ = 0, the limit of infinite sample size [Strong et al., 1998]. We report the extrapolated value as our final estimate. The resulting information value 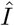 has the units of bits. To convert 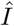 to bits per second, we multiply by it by *v*_*max*_*/*0.03.

### Computing Mutual Information from Simulations

For Figure 2 we plotted the Shannon Mutual Information (*I*) under various settings of the population parameters, for various values of the population size *N*. To compute 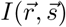 we use the following formulation of *I*:

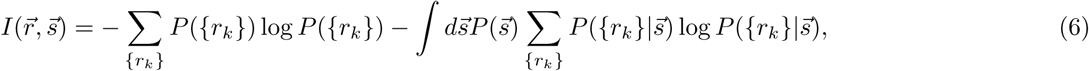

where 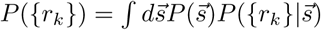. Expectation with respect to 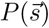 was approximated by averaging over *N*_*s*_ = 5000 samples drawn from 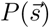, which was the uniform distribution on the two dimensional unit circle. Because both the information-preserving population vector 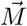 and the standard population vector 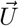 are discrete random variables like 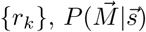 and 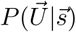 are computed by pooling 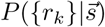 across all *{r*_*k*_*}* that map to the same value of 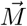 or *_ 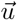*. These discrete mappings are precomputed. Once we have computed 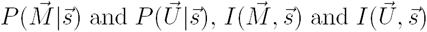 are computed in the same way as 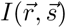.

The simulated neural populations had the following parameters in Figure 2. In panels A and B *φ*_*i*_ = 0 for all *i*. In panel A the *β* values are uniformly distributed on a log10 scale between 0.1 and 10, and the *α* values are set so that the peak firing rate equals 0.8 for all neurons (see Eq. (S5) below). In panel B all *β* are set equal to 1 and the *α* values are set so that the peak firing rate varies uniformly between 0.4 and 0.8. In panels C and D the population is divided into subpopulations of equal size with preferred orientations at *±*45 degrees. In panel C, for both subpopulations, the *β* and *α* values are set as in panel A. In panel D the same *α* and *β* values are used as in panel C though the peak firing rate may differ from 0.8 due to the presence of interneuronal coupling induced by noise correlations. The noise correlations parameters were set according to Eq. (S25) below.

In the case of neural populations tuned to the same stimulus feature, we also computed information while binning the response statistics. The standard population vector reduces in this case to the population count variable *U*_count_. The information-preserving population vector also becomes a scalar variable *M*_count_ = Σ *r*_*i*_*β*_*i*_. To compute the binned versions of these quantities *U*_bin_ and *M*_bin_, we divided the support of either *M*_count_ or *U*_count_ variables into 15 equal sized bins. We then computed the mappings assigning values of *{r*_*K*_*}* to values of *M*_bin_ and *U*_bin_, using the mappings from *{r*_*K*_*}* to *M*_count_ and *U*_count_ as an intermediate step. The results are included in Figure 2 as dotted lines. They indicate that even a small number of bins is sufficient to capture essentially all the information provided by the responses of a neural population.

### Text S1 Linear-Nonlinear Model and Orientation Tuning

We begin by modeling neural responses from individual neurons as a binary variable *r* taking a value 1 when the neuron produces a spike and 0 otherwise. To account for response saturation and rectification, we model the probability of a spike (*r* = 1) as a saturating function of the stimulus projection onto the neuron’s receptive field 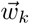 Specifically, we choose the logistic function in order to take advantage of the properties of exponential families described below.

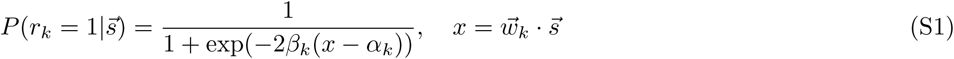

Here, vector 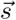 represents the current stimulus, 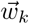 represents the preferred stimulus or receptive field (RF) of the *k*th neuron, and *x* is the component of the stimulus along the receptive field. The parameters *α*_*k*_ and *β*_*k*_ describe, respectively, the midpoint and slope of the logistic function (Figure 1a). As a matter of notation neurons will be indexed by the letters *i, j,* and *k* and dimensions of the stimulus or neural receptive fields will be indexed by *a, b, c,* and *d*. The RF can be thought of as a pattern of unit contrast that, if presented, would elicit the strongest response from the neuron. Both 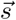 and 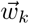 are *D*-dimensional vectors and 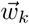 is assumed to be normalized. We note that if a neuron with an orientation sensitive receptive field, such as the one shown in the inset of Figure 1a, were probed by stimuli of oriented gratings with fixed contrast level then the nonlinearity described in (S1) would yield a typical tuning curve around the preferred orientation (Fig. 1B). Instead of considering such a high dimensional receptive field we work with a simplified model of orientation tuning in order to provide a clearer link between the parameters *α*_*k*_, *β*_*k*_ and the shape of the orientation tuning curve. Specifically, for a neuron with preferred orientation *φ*_*k*_ we define RF as 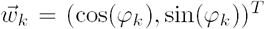, while stimuli are described by 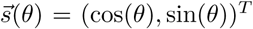. Thus, in the framework of the linear-nonlinear model, the probability to observe a spike is given by:

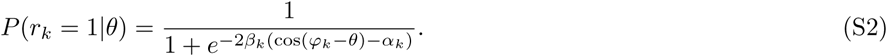

The maximal spike rate is achieved for *θ* = *φ* _*k*_, which is given by:

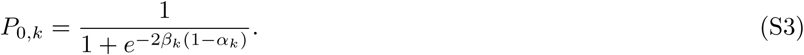

The width of the orientation tuning curve, which we define as the inverse of the second derivative of the logarithm of the tuning curve is:

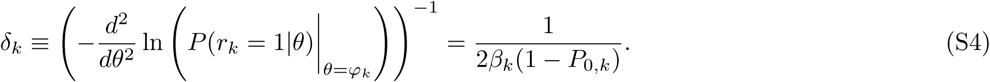

We can invert the above equations to express *α*_*k*_ and *β*_*k*_ in terms of *P*_0,*k*_ and *δ*_*k*_:

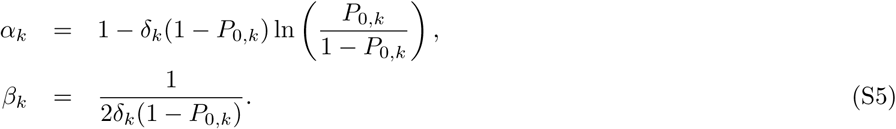

Thus, there is a one-to-one correspondence between parameters of tuning curves and nonlineary of the LN model.

### Text S2 Information-Preserving Population Vector

### Population response probability is an Exponential Family

We start by describing the case where neural responses are conditionally independent given the stimulus; the generalization to the case of correlated neural variability will be discussed in Text S3. Further, we will begin our arguments by considering the responses of a population of neurons at one time point, and in a time window small enough for the responses of individual neurons to be binary. At the end of Text S2 we will discuss how the results can be generalized to tackle the case of longer time windows where neurons produce multiple spikes.

It is helpful to re-write the response function of an individual neuron (S1) in an exponential form:

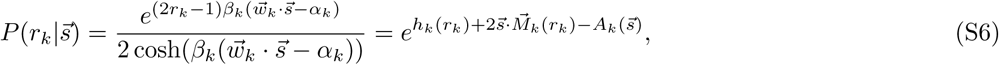

where we have defined the functions 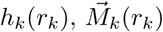, and 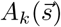 for notational convenience:

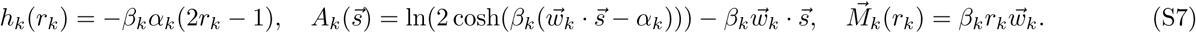

For conditionally independent responses, the probability to observe a response pattern *r*_1_, *r*_2_, *…, r*_*N*_ across *N* neurons to stimulus 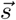 is:

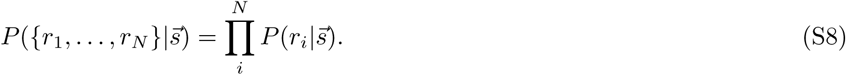

Using Eq. (S6), this probability distribution can also be written in the exponential form:

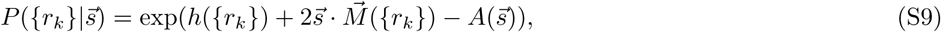

where the functions 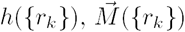 and 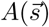 are the summations of the corresponding individual neuron functions:

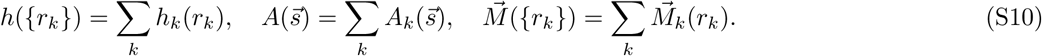

The vector 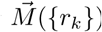 is the information-preserving population vector whose expression we explicitly write out because of its importance:

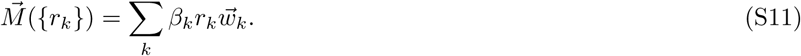

It provides a mapping from a set of neural responses {0, 1}^*N*^ to a finite subset of ℝ^*d*^. The important conclusion is that the population response model (S9) forms an exponential family with natural parameter 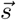 and sufficient statistic [Wainwright and Jordan, 2008] 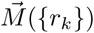 that corresponds to the information-preserving population vector. We will see in Sec. Text S3 that the population response model with correlated variability across neurons also forms an exponential family As a matter of terminology, a single exponential family is considered to have fixed values of *{α*_*k*_*}, {β*_*k*_*}*, and 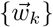 with different members of the family indexed by different stimulus values of 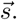

### Sufficient statistics preserve information

An important result of 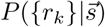 being an exponential family is that the mutual information is preserved by the sufficient statistic [Cover and Thomas, 2012], which in our case is the information-preserving population vector *M* from the main text. That is, our goal is to show that:

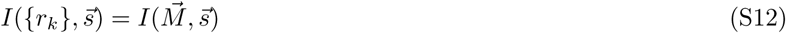

To show this directly in our case we first define the “density-of-states” function 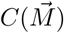 as the sum of *e*^*h*(*{r*^*k}*) over all {*r*_*k*_} that map to the same value of 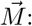

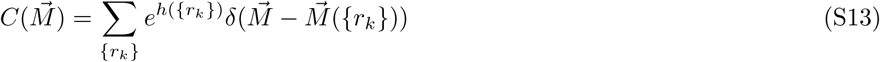

The conditional and marginal distribution of 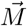 can be expressed in terms of 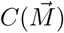, without direct reference to {*r*_*k*_}:

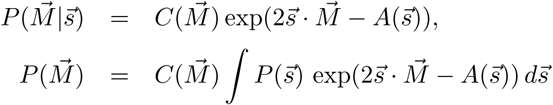

We note the relationships between 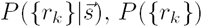 and 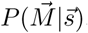, 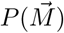 respectively:

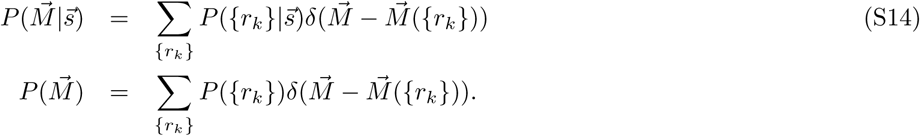

We now have the following important identity:

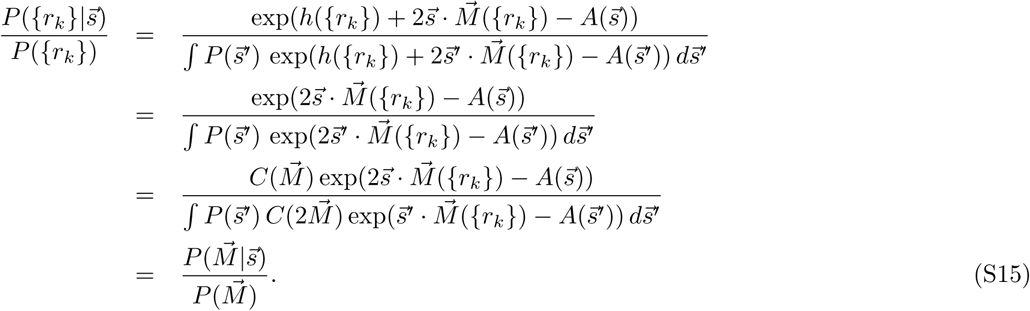

This last equality applied to Eq. (S12) for the mutual information yields:

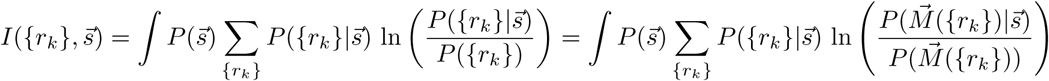

This in turn yields that

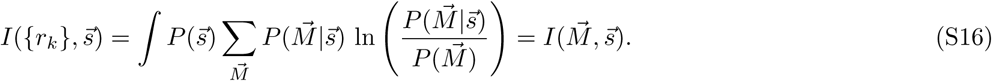

Another corollary of (S15) is that the posterior distribution of 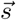 given {*r*_*k*_} depends only on 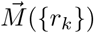:

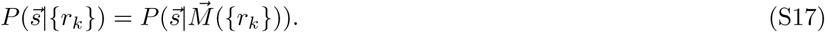

Therefore, a Bayes optimal decoder needs only to carry out the weighted summation rather than keep track of which response (out of 2^*N*^ possible) was observed. Similar sufficiency properties are known for Gaussian *r*_*k*_ as well as binary population models with independent and identically distributed neurons [Wainwright and Jordan, 2008, Ma et al., 2006]. The derivation provides the first demonstration, to our knowledge, for a sufficient statistics for a population model for binary neurons that are neither independent (see also Text S3 below) nor identically distributed and which has dimension *D* independent of population size (*D* is the stimulus dimension).

### Cumulants of the information-preserving population vector 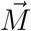

The mean value of the information-preserving population vector varies smoothly with the stimulus as we illustrate in Figure S1. To show this analytically, we provide in this sub-section analytic expressions for the first two cumulants of 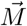 as a function of 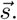 Since 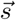 is the natural parameter of the class of models we consider, the cumulants of 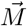 can be computed by taking derivatives of different orders of the log-partition function 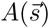 with respect to 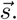. In particular the mean and covariance are the gradient and Hessian respectively:

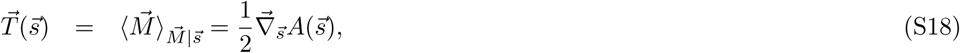

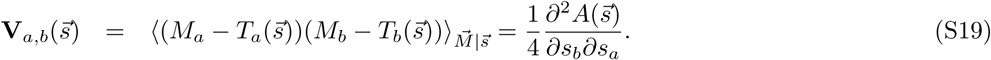

Since covariance matrices are always positive semi-definite we see here that 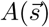 is a convex function. We also note that 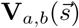 is the Jacobian of 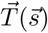, as well as the Fisher information matrix of 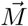 with regards to 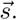 When the neurons are conditionally independent, 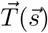 and 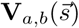 take simple forms:

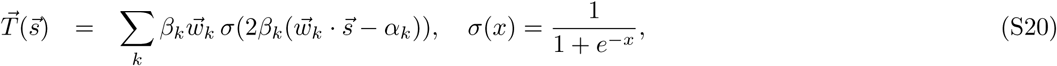

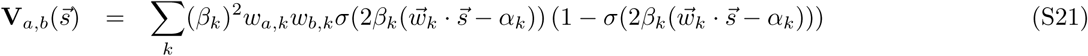

Because 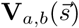 is a continuous mapping we see here that 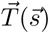 is a smooth function of 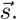 It is sometimes the case that stimulus parameter is embedded in a higher dimensional space as the natural parameter 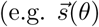 has higher dimensionality then *θ*, a single parameter). In this case the family is said to be *curved* with respect to *θ*. We note that the information-preserving property also holds for curved exponential families. However, when calculating the cumulants, it is necessary to take gradients with respect to 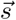 and not *θ*.

### Analysis of non-binary neural responses

We now discuss how these analyses can be generalized to non-binary neural responses that may appear over longer time windows. The time window size *T* still needs to be constrained by the stimulus dynamics to ensure that the stimulus does not change appreciably during the response time window. Nevertheless, for natural visual stimuli it would not be uncommon for stimuli to be approximately constant over the time period of *∼* 30 msec, given the predominance of low temporal frequencies in natural stimuli [Simoncelli and Olshausen, 2001]. Over this time window the responses of visual cortical neurons, for example, would commonly produce more than one spike. How should we treat such multiple responses? Formally, we can split the time window of interest *T* into smaller bins of width Δ*T* of such duration that the neural response can only be binary (e.g. *∼* 1 msec) when considered in these Δ*T* time intervals. The maximal number of spikes that a neuron can produce is then *N*_*t*_ = *T/*Δ*T*. We can model responses of this neuron that as a set of *N*_*t*_ binary neurons with identical LN parameters. Therefore, applying the same mathematical arguments as above, one observes that the responses of these auxiliary neurons can be simply averaged without incurring information loss. (These summed responses will follow a binomial distribution.) Returning to the population of *N* neurons with different RFs and non-binary responses, we can expand this population to size *N*_*t*_ *· N* where each neuron from the original population is represented by a subpopulation of *N*_*t*_ neurons, with the same RF and *β* factors as the original neuron, and whose responses can therefore be averaged. This analysis indicates that the responses of non-binary neurons can be analyzed by fitting the neural responses to stimuli with a logistic function scaled by a constant *R*_max_. The inverse of the scaling constant *R*_max_ yields the time window duration over which the responses of this neuron can be considered binary. The expression (S11) for the information preserving population vector remains unchanged, 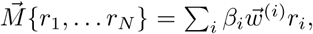 except now *r*_*i*_ are no longer binary variables and instead represent the number of spikes produced by *i*th neuron during time interval *T*. Furthermore, because the response averaging holds in the presence of noise correlations, as long as they are stimulus-independent (see Text S3 below), the averaging across time will be valid even when responses across different time bins are not independent, again as long as these correlations across time bins do not depend on the stimulus.

### Text S3 Taking into account correlated variability across neurons

We now expand the population response model to allow for the presence of correlated variability among neurons observed under repeated stimulus presentations. A standard way to include such pairwise “noise” correlations between neurons is with the following probability response distribution [Granot-Atedgi et al., 2013]:

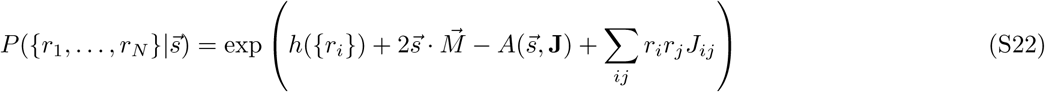

The new term Σ_*ij*_ *r*_*i*_*r*_*j*_*J*_*ij*_ that describes noise couplings between neurons does not depend on the stimulus, and so can be incorporated into *h*({*r*_*k*_}):

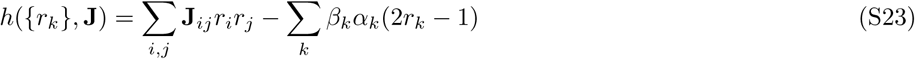

The joint distribution on population responses is again an exponential family:

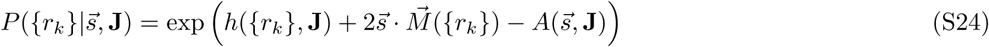

Importantly, the vector of sufficient statistics 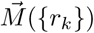 is the same as before and still preserves information in neural responses. Thus, the strategy for reading out the activity does not need to be modified in the presence of correlations. The normalization factor 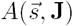 is now defined as a stimulus dependent normalizing term but in general lacks a closed-form expression similar to 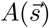 from Eq. (S10).

In the simulations in Figure 2D, we used coupling coefficients parameterized by {*φ*} as follows:

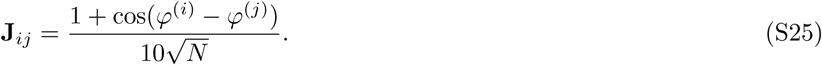

It is worth nothing that the framework remains valid in the presence of noise correlations that depend on differences in the stimulus selectivity between neuronal pairs. We note that for non-zero **J**, computing 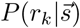 requires marginalizing over the states of all other neurons in the population and will in general differ from the logistic response function Eq. (S1). This observation demonstrates that it is not the logistic form Eq. (S1) that guarantees 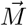 preserves information but rather that the population’s response distribution is coupled to 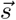 only through 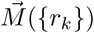. Furthermore, since the expressions relating the cumulants of 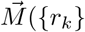 to gradients of 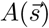 are still valid in this case and 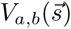 remains positive semi-definite by construction, the mapping described by the expected value 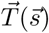 of the information-preserving population vector remains a smooth function of stimuli 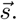

### Text S4 Stimulus Decoding from Population Vectors

We now describe convergence properties of the information-preserving population vector in the limit of large neural populations. Here, the response function of individual neurons does not have be to a logistic function for most of the important properties to hold. Therefore, we write this response function from Eq. (S1) more generally in terms of a smooth monotonic function *g*:

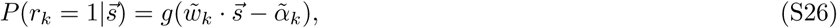

where we introduced a new notation 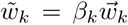 for the RF scaled by its amplitude *β*_*k*_. The thresholds have also been correspondingly scaled 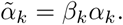

We will normalize both the information-preserving and the standard population vectors by the number of neurons *N* in the population

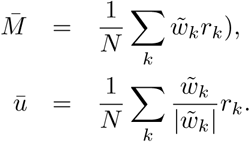

By the weak law of large numbers both 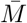 and 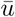 converge in probability to their expected value as *N* grows large:

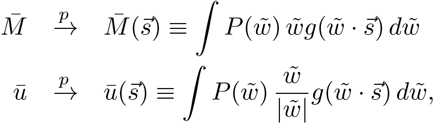

where 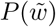 describes the distribution of (scaled) RF components. We now show that when 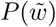 is described by a multivariate Gaussian distribution, the information-preserving vector will produce unbiased stimulus estimates, whereas biases will persist in reconstructions based on the standard population vector. For clarity of the presentation, we first consider the case where RF components have zero mean and that all neurons have the same scaled thresholds 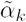 Later we will show how solutions generalize to the case of unequal thresholds and nonzero RF components.

Just like in the analysis of STA convergence properties [Paninski, 2003, Sharpee et al., 2004], we can use Stein’s lemma for Gaussian 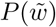 [Stein, 1981] that expresses averages of RF components weighted by the nonlinear function *g* of these components as the product of correlations between components and the average of the gradient of *g*:

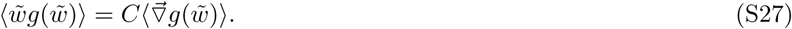

Here ⟨ *·* ⟩ denotes expectation with respect to 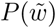 and 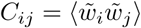 is the covariance of RF components across the population. Stein’s lemma applies as long as *g* is a smooth function. [Technically, 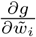 should exist almost everywhere and 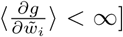. Applying the Stein’s lemma to Eq. (S27) for the information-preserving population vector one finds that:

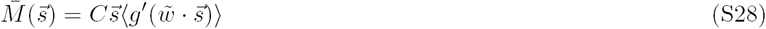

In these equations, the average 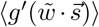 describes the compressive nonlinearity of the kind shown in Fig. S1. The important conclusion from Eq. (S28) is that the information-preserving population vector is aligned with 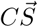. The stimulus direction can therefore be determined by multiplying the information-preserving population vector by the inverse covariance matrix *C*^−1^. This procedure is completely analogous to the one used to reconstruct the RF from the STA in the presence of stimulus correlations [Sharpee et al., 2004, Ringach et al., 2002].

One finds a very different answer when applying Stein’s lemma to the standard population vector. In this case:

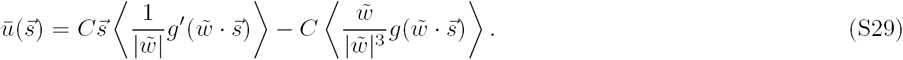

Here, the average in the second term will not be aligned with the stimulus 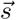 unless the input distribution has spherical symmetry [Chichilnisky, 2001, Paninski, 2003]. Thus, the right hand side is not aligned with 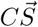 This indicates that that decoding based on the standard population vector does not readily produce an estimate of the stimulus, even in the limit of large neural populations.

Standard numerical issues might arise when computing the inverse of the covariance matrix *C* if RFs primarily sample one portion of the stimulus space. These issues have been discussed in detail in the context of RF estimation [Sharpee et al., 2008, Sharpee, 2013, Ringach et al., 2002, Theunissen et al., 2000]. In Figure 4 we plot, the correlation between stimuli 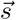 and *C*^−1^*M*. However, one can also follow [Simmons et al., 2013] to bypass these issues by analyzing the correlation between 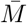 and 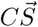:

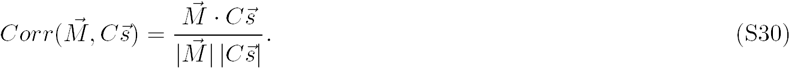

We now return to consider generalizations to the case where RF components have non-zero mean and thresholds vary among neurons. To compensate for the non-zero mean, one can subtract from the information-preserving population vector a vector of 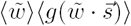, the average *RF* scaled by the population firing rate in response to 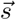 This linear procedure can be achieved in the brain in the balanced regime [van Vreeswijk and Sompolinsky, 1996] where excitatory and inhibitory inputs are balanced on average together with homeostatic scaling of synaptic inputs.

The case of variable thresholds is treated by considering the average nonlinear function

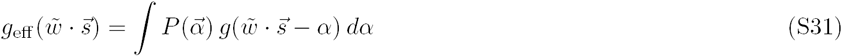

The above results remain valid as long as *α* are distributed independently of 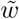

Finally, the distribution of RF components might not be purely Gaussian. Such deviations will cause systematic distortions in the mapping (Fig. S1). As long as these distortions are indeed systematic, they can be learned and compensated for. Such learning also needs to happen every time the RF changes following adaptation to changes in the stimulus statistics. One possible approach for computing the expected deviations of the estimator Eq. (S27) for weakly non-Gaussian stimuli can be found in Appendix A of [Sharpee et al., 2004].

The question of what sets of receptive fields might provide maximal information about natural scenes and allow for their accurate reconstruction represents an active area of research [Olshausen and Field, 1996, Olshausen and Field, 1997, Henniges et al., 2010, Bornschein et al., 2013]. Here we pursued a separate question of how to decode neural responses based on a fixed set of receptive field while minimizing information loss. Finding optimal receptive field parameters is an important task for future research, with results that will likely differ for linear-nonlinear or quadratic decoding.

### Receptive field decorrelation by recurrent networks with divisive normalization

According to Eq. (S28), the information-preserving population vector yields an estimate of the stimulus 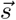 multiplied by the covariance matrix *C* of neural RFs in the population. This systematic shift can be compensated for by “decorrelating” RFs to applying such transformations that the resultant covariance matrix becomes proportional to a unit matrix. We now discuss how divisive normalization in a recurrent network can approximate this operation.

In the divisive normalization model [Carandini and Heeger, 2012] neural responses depend on the activation of other neurons in the network as follows:

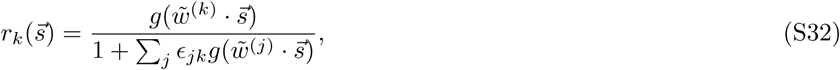

where 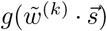 describes the activation function of the *k*th neuron without taking into account recurrent connections. The activation function *g*(*x*) is again a smooth, monotonically increasing function. The parameters *ϵ* _*jk*_ describe the strength of the recurrent connection from neuron *j* to neuron *k*. In general they are not symmetric, 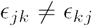. We let 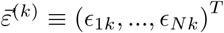 denote the vector of incoming connections to neuron *k*.

We consider effective receptive fields 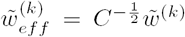 the Cholesky decomposition of *C*^−1^), so that the covariance matrix of 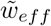 across the population is the *D*-dimensional identity matrix **I**. We seek to find a settinVthat 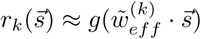 as close as possible:

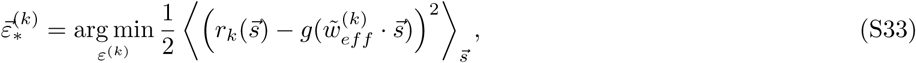

where the stimuli are drawn from a distribution 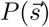. We note that the optimization in (S33) can be carried out independently for each *k*. We performed an example computation with the following parameters: *D* = 2, *N* = 500, 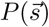 was a white noise gaussian that we drew 1,000 samples from. Rfs 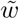 were drawn from a zero mean Gaussian distribution with the following covariance matrix:

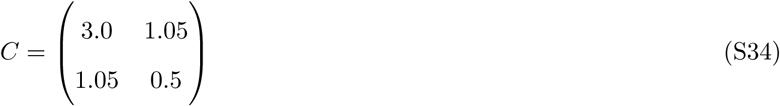

The optimization in (S33) was computed using L-BFGS algorithm, and *∊*_*jk*_ was initialized as cos(*θ*_*jk*_)*/N* where *θ*_*jk*_ is the angle between 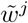 and 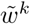. This initialization was chosen to ensure that no numerical overflow occurred in the evalation of 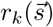. After finding 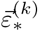 for all *k*, we computed the pearson correlation between *ϵ* _*jk*_ and cos(*θ*_*jk*_). For this example we found a correlation of 0.02 with p-value *p* = 4.2 *×* 10^−23^, indicating a weak but statistically significant correlation between the strength of recurrent connections and receptive field overlaps, as is observed experimentally [Yoshimura and Callaway, 2005, Yoshimura et al., 2005].

### Text S5 Quadratic Expansion of the Input Space

To account for non-monotonic spike probabilities as well as the dependence on multiple stimulus components, one can follow the framework of minimal models [Fitzgerald et al., 2011] to expand the stimulus by including all pairwise products between stimulus components. If the original stimulus has *D* components, the expanded stimulus will have *D* + *D*^2^ components of the following form:

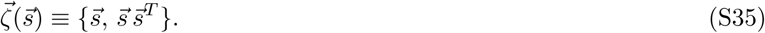

The neural response probability

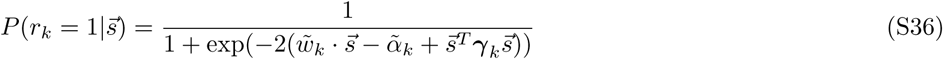

can be compactly written as

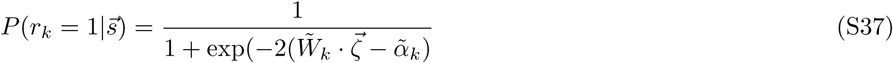

in terms of the RF 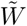 of D + D^2^ dimensions in the expanded input space. Thus, the information-preserving properties analyzed in Text S2 and decoding properties discussed in Text S4 will hold. We note that the quadratic matrix *γ* can have both positive and negative eigenvalues to describe both excitatory and suppressive stimulus dimensions [Rust et al., 2005].

The information-preserving population vector will now also have *D* + *D*^2^ components:

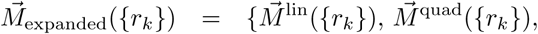

where

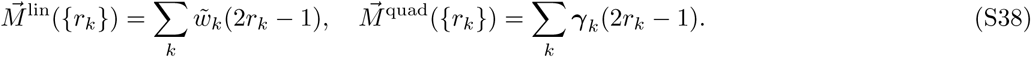

If both 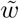 and ***γ*** are normally distributed, then, following the arguments from Text S4, the information-preserving population vector in the expanded space will be aligned with *C*_expanded_*ζ*. Here, *C*_expanded_ is the covariance matrix of the *D* + *D*^2^ dimensional vector 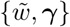. On a particular trial, 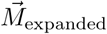 may produce a direction *ζ* that does not satisfy the constraint of Eq. (S35) associated with quadratic expansion of the stimulus. However, we can search for a value of *ζ*_est_ that satisfies this constraint and is most consistent with 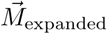. This will in turn produce a stimulus estimate in the original input space. To find such a pattern, we note that 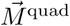 is a symmetric matrix. The best rank one approximation of this matrix, according to the Eckart–Young–Mirsky theorem, is 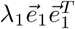 where *λ*_1_ is the largest (in absolute value) eigenvalue of 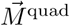 and 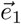 its associated eigenvector. We set 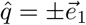 and resolve the ambiguity in sign by requiring consistency with the estimate derived from 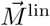.

The power iteration algorithm can be used to find *_e*_1_. For our purposes, we initialize 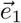 randomly and apply the following iteration:

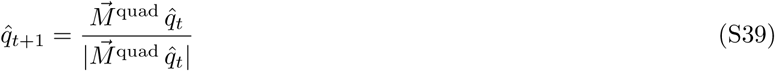

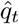 will eventually converge to 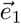. We note that the iteration in (S39) involves a sequential application of matrix multiplication by 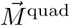 (recurrent processing), followed by normalization of the resultant vector (gain normalization). The vector 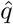 provides the best approximation of *M*_expanded_ that can be constructed from *D* dimensional vectors as 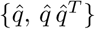.

For the decoding of V1 data, we took 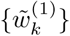 and 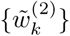 to be the first and second receptive field components of [Sharpee et al., 2006]. The quadratic kernel *γ*_*k*_ is computed as 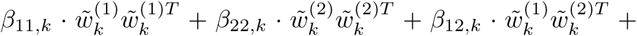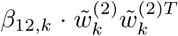, where coefficients *β*_11_, *β*_12_ and *β*_22_ are defined in Eq. (2). In this case it can be shown that under the assumption that RF components 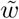 are described by a Gaussian distribution with zero mean, the covariance matrix in the expanded space allows a decomposition such that

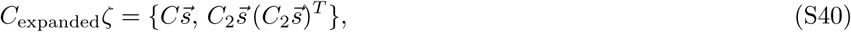

where 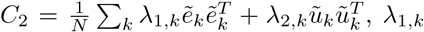 and *λ*_2,*k*_ are the first and second eigenvalues of the quadratic kernel for *k*th neuron, and 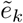 and 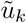 are their corresponding eigenvectors. Specifically, 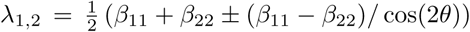 and *θ* = arctan(2*β*_12_*/*(*β*_22_ *- β*_11_))*/*2. Thus, we did not need to compute the full covariance matrix in the expanded space. Comparing Eq. (S40) and the definition of 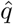, we can find the stimulus direction based on quadratic decoder by applying 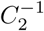 to 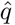. Alternatively, to avoid inversion of a poorly conditioned *C* matrix, we compute, just as in Text S4, the vector correlation between 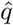 and 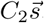.

**Figure S1:**
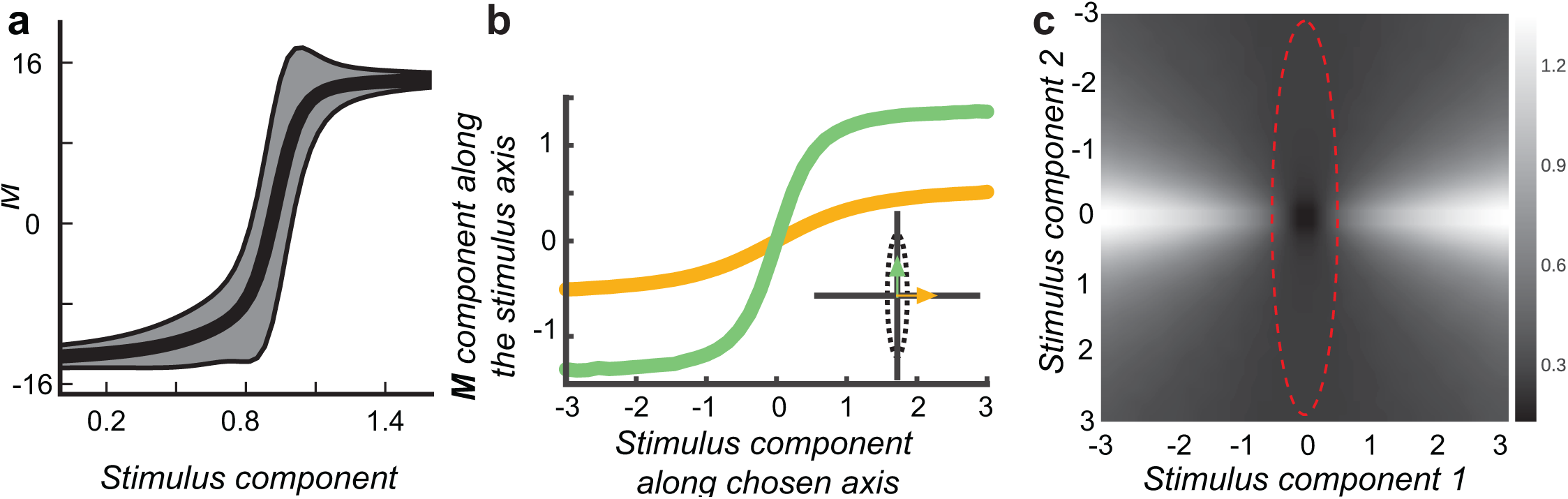
(**A**) The expected value of the information-preserving spike count varies smoothly as a function of the stimulus component along the RF. Gray shading represents one standard deviation around the mean (black line). (**B**) Illustration of the compressive nonlinearity in a population tuned to different input features. The curves relate the magnitude of stimuli along one of two axes in the input space (green or yellow) to the magnitude of the information-preserving population vector along these axes. Green/yellow curves are for directions of maximal/minimal variance of RF components. (**C**) Map of the compressive nonlinearity showing the magnitude of the information-preserving population vector multiplied by *C*^−1^ (to correct for differences in RF distribution) for different stimuli, see also Text S4. The red dotted line shows the standard deviation of the RF distribution.

